# Yttrium-iron garnet magnetometer in MEG: Advance towards multi-channel arrays

**DOI:** 10.1101/2022.07.11.499607

**Authors:** E. Skidchenko, A. Butorina, M. Ostras, P. Vetoshko, A. Kuzmichev, N. Yavich, M. Malovichko, N. Koshev

## Abstract

Recently, a new kind of sensor applicable in magnetoencephalography (MEG) has been presented: a solid-state yttrium-iron garnet magnetometer (YIGM). The feasibility of YIGM was proved in alpha-rhythm registration experiment. In the current paper, we propose the analysis of lead-field matrices for different possible YIGM multichannel on-scalp sensor layouts with respect to the information theory. We use real noise level of the new sensor to compute signal power, signal-to-noise ration (SNR) and total information capacity, and compare them with corresponding metrics that can be obtained with well-established MEG systems: based on superconducting quantum interference devices (SQUIDs) and optically-pumped magnetometers (OPMs). This simulation study is aimed to shed some light on the direction for further development of YIGM sensor, namely creation of multi-channel YIG-MEG.

## 1 Introduction

### 1.1 History

Magnetoencephalography (MEG) is one of the modalities for neuroimaging that allow to visualize and research the electrical activity of the brain. The MEG technique is based on measurement of the magnetic induction produced by cortical currents, with an array of sensors (magnetometers) located outside a head. Due to the fact that biological tissues are transparent to magnetic fields, MEG dramatically exceeds the electroencephalography (EEG) technique in terms of spatial resolution and frequency sensitivity. On the other hand, magnetic field is a direct consequence of electric current, which allows MEG to outperform methods based on magnetic resonance imaging (MRI) in terms of temporal resolution. Summarizing all above, the MEG technique combines high temporal and high spatial resolutions as well as good sensitivity to high-frequency brain currents, which makes MEG the most powerful among non-invasive neuroimaging tools.

The main technological difficulty of the MEG caused by weakness of magnetic fields inside the region of measurements. The primary cortical currents are of the *nA*-order. Taking into account the fast decay of magnetic field with distance from a source, in the measurement region it has the *fT*-order. This requires application of extremely sensitive magnetic sensors and magnetic screens (magnetically shielded room, or MSR) with high screening coefficient.

For the first time MEG was demonstrated by D.Cohen *et al*. [11] in 1971, proposing to use superconducting quantum interference devices (SQUIDs) to measure brain-induced magnetic fields. Since that time, for almost half a century SQUIDs were the only sensors capable of registering MEG signals. In different SQUID systems, the scalp-to-sensor distance varies within 2-3 cm, but it is no less than 2 cm. The sensitivity of the SQUID systems reaches 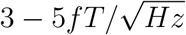 [21]. However, SQUID sensors have several significant drawbacks related to demand of cooling them with liquid helium to maintain the superconductivity phenomenon. SQUID-based devices are very expensive and bulky. Moreover, SQUID helmets are not adaptable to different head shapes and sizes, which leads to appearance of additional errors in measurements. Bulkiness of the SQUID-based MEG-systems require to use of big and heavy MSRs that are expensive itself. All these disadvantages explain respectively low popularity of MEG despite its high capacity.

Recent technological advances in the field of high temperature superconductivity (high-Tc SQUID) as well as in the field of atomic optically-pumped magnetometers (OPMs) may partially lift some limitations of conventional SQUID-based MEG technique, sufficiently spreading the popularity of MEG.

MEG systems based on high-Tc SQUID sensors [14] are bulky too, however, the technology is quite promising. They operate at liquid nitrogen temperature instead of helium, having sensitivity close to that of conventional low-Tc SQUIDs. This fact makes sensors much cheaper in terms of both CAPEX and OPEX. The high-Tc SQUIDs may also be placed much closer to a scalp (about 1-2 mm), advantages of which are demonstrated, e.g. in works [3,4]. In the last decade, the MEG market met OPMs. Demonstrated to be feasible for MEG purposes in 2010 [34], these sensors became currently the main candidate for being the future of MEG, which is supported by a big number of OPM-driven research in the field. OPMs do not require cooling to cryogenic temperatures for operation and can be assembled into flexible arrays around the head, being adapted to any head size and shape and providing respectively small scalp-to-sensor distance (about 6-6.5 mm). All these facts lead to appearance of a number of laboratory and commercial OP-MEG systems ([5–8, 25], etc.). Indeed, appearance of OPMs breathed new life into MEG, and the OP-MEG studies are being continued nowadays [18, 24, 29, 44]. Nevertheless, OPMs, in their turn, have disadvantages of narrow dynamic range (±1.5−5 *nT* [2,9,37]), heating problems and limited lifetime of the sensors.

However, a really big step forward in MEG evolution was done recently, in 2021, when the new type of highly-sensitive sensor had been presented and demonstrated to be feasible in terms of MEG [23]. The sensor is based on yttrium-iron garnet thin films and inherits the principle of work of a flux-gate magnetometer. Yttrium-iron garnet magnetometer (YIGM) is a solid-state sensor and unlike SQUIDs and OPMs operates at room temperatures, is durable and shows comparatively wide dynamic range (10 *µ*T [41]). Advantages of the sensor allow to assume appearance of a new promising YIG-MEG technique in the nearest future.

### 1.2 Motivation

Regardless of the nature of novel sensor types, development of MEG system on the base of them demands a big amount of research work in the field of effectiveness and information capacity of the system under development. The task is non-trivial and previously has been considered by researchers all over the world. It requires application of mathematical modeling and information theory.

The amount of information being gathered by an array of magnetometers depends on a set of factors: sensors’ shape and dimensions, scalp-to-sensor distance as well as orientation of sensitive axes in magnetometers with respect to orientation of sources, particular features of particular cognitive experiments, sensitivity/intrinsic noise of sensors in array, positions of the sensors in array, etc. Some magnetometers are designed to measure only radial (normal) component of the magnetic induction, while the others provide measurements of two or even three orthogonal components [10], commonly, with lesser sensitivity [2].

The current work is dedicated to the research of potential of multi-channel YIGM sensor arrays in terms of MEG. This research is motivated by completely different nature of the sensor and specifics of its geometry. The zero-generation of the YIGM is quite big in dimension (38×38×3 *mm*^3^) and noisy - the level of intrinsic noise is about of 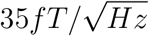 [23]. The work [41] shows the possibility of decrease intrinsic noise down to 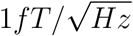 (theoretical limit) and size with increase in pumping frequency. However, decrease in both size and intrinsic noise demands a big amount of engineering work of different kind. Thus, we need to define the optimal development strategy for new generations of YIGM sensors.

Being flat by design, the YIGM may be placed in different ways in order to register either normal (radial) or tangential component of magnetic induction with respect to head (see Section 4.1). Since registration of tangential component means closer distance from sensor to scalp compared to registration of normal component (see [23]), it is reasonable to investigate and compare the quality and amount of information that can be gained in both these ways. The information quality and capacity have been previously discussed in a set of research. Iivanainen *et al*. [21] shows, e.g. the greater impact of volumetric Ohmic currents on the tangential component of the magnetic induction compared to that on radial component. The works [17] and [20] show, however, absence of theoretical grounds of this phenomenon, while [22] shows insignificance of extra computational complexity of using purely tangential components with respect to inverse problem solution.

Dependence of the information capacity on scalp-to-sensor distance has been explored by different research teams due to recent technological advances in the field of high-Tc SQUIDs and OPMs. For example, two experimental research [3, 14] compare the responses registered by conventional low-Tc SQUID-MEG systems with those by high-Tc magnetometers placed in several millimeters from scalp, showing considerable increase of signal-to-noise ratio for the latter. Analysis of the lead-field matrix [21, 33], and inverse problem solution [31, 38] also show significance of sensors’ placement closer to head, estimating the signal-to-noise, total information capacity, and spatial resolution of the inverse problem solution.

In this paper, we will explore lead-field matrices for different possible YIGM multichannel layouts, analyse the metrics according to the information theory and compare them with the metrics for corresponding layouts for arrays of SQUID and OPM sensors.

## 2 Theory

This section is dedicated to brief physical and mathematical descriptions of the phenomena and techniques underlying the MEG method.

### 2.1 Governing equations

In order to model electric and magnetic fields produced by cortical currents, the head is considered to be a closed piecewise homogeneous volume conductor ([15, 16, 32], etc.) Ω ∈ ℝ^3^ with boundary ∂Ω. The interfaces inside this volume are indexed with index *s* = 0, …, *S* and denoted by ∂Ω_*s*_. The electric current density can be presented in two parts:

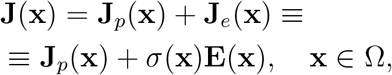

where **J**_*p*_(**x**) represents the primary currents occurring in dendrites, and **J**_*e*_(**x**) = *σ*(**x**)**E**(**x**) is the Ohmic volume currents driven by non-zero electric field **E** in a volume conductor.

Due to comparably low frequencies of the sources, the quasi-stationary approximation can be applied, which allows to reduce the Maxwell equations to two Poisson-like elliptic governing equations for electric and magnetic fields [12, 30]:

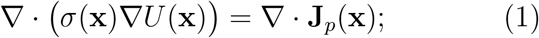

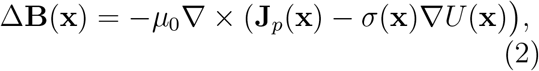

where *U*(**x**), **x** ∈ Ω is the electric potential, *σ*(**x**) represents the tissue conductivity at the point **x** ∈ Ω, and *µ*_0_ is the vacuum permeability.

Finally, after integration of the equation for **B** and its transformation due to integral identities [15], we end up with the following formula for the magnetic induction:

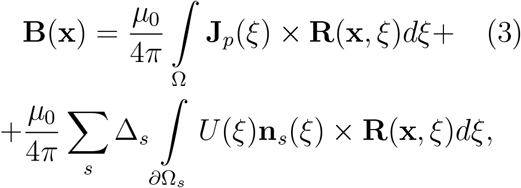

where **n**_*s*_(*ξ*) is the normal to the surface interface ∂Ω_*s*_ at the point *ξ* ∈ ∂Ω_*s*_, Δ_*s*_ is the difference in tissue conductivity between volumes divided by the interface ∂Ω_*s*_, and **R**(**x**, *ξ*) is the vector function introduced for brevity:

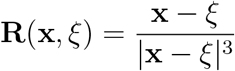

Thus, the magnetic induction can be calculated via simple integration after the computation of the electric potential using the equation (1) with proper boundary conditions. The output of the sensor is commonly obtained by integration of the magnetic induction over the sensitive volume of the sensor under consideration (see Subsection 4.2).

### 2.2 Lead-field matrix

Natural discretization of the equation (3) allows to obtain the sensor outputs for *N*_*s*_ sensors in the simple linear approximation [16, 35]:

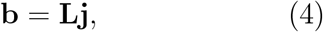

where the vector 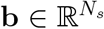 represents scalar outputs of *N*_*s*_ sensors, 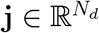 is the vector representing current dipoles at *N*_*d*_ discrete points on the cortex, and 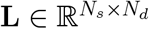 is the lead-field matrix obtained with MRI-based head geometry, proper conductivities and taking into account the integration over the sensors’ working body and geometry of its sensitive axes. The rows of the lead-field matrix represent the outputs of sensors for unit sources stored in columns: *L*_*ji*_ is the output of *j*^*th*^ sensor if only *i*^*th*^ unit source is active.

Construction of the lead-field matrix **L** is the most common way to solve the MEG forward problem. The number of source defines its position, and the overall number of dipoles *N*_*d*_ depends on the desirable accuracy of source localization.

On the base of **L** measurements one may compute the estimation of the underlying sources **j**, i.e. solve the *MEG Inverse Problem*. The inverse problem is ill-posed according to Hadamard due to its solution being unstable, and underdetermined due to *N*_*s*_ < *N*_*d*_. There are different ways to solve this problem, the main ones are beamformer [19], MNE [16], LORETA [28], MCE [39]. In the current research, we use the most conventional way to do it: minimum norm estimation, or MNE. This method is close to Tikhonov’s regularization and implemented almost in every library or package for solution of MEG inverse problems. In the paradigm of MNE, the source estimate can be presented as follows [35]:

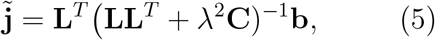

where *λ* > 0 is the regularization parameter limiting the *l*_2_-norm of the solution, and **C** is the noise covariance matrix. Due to [26], the regularization parameter may be computed using the estimated signal-to-noise ratio (SNR) via the following expression:

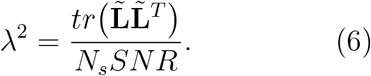

Here the 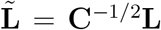 denotes lead-field matrix after whitening transformation.

## 3 Methods

The current research was inspired by works [33] and [21]. The first work proposes adaptation of Shannon’s channel capacity to MEG sensor arrays, forming the metric called ”information capacity”, while the second one considers a lot of metrics, including topography power, SNR of source, relative signal power, relative SNR, analysis of the point spread functions, and total information as well.

For the first research related to YIGM arrays information quality, we compute relative signal power, relative SNR, and total information capacity. For convenience, we briefly describe these metrics below in the current section.

All the metrics presented below take into account the sensor’s noise covariance matrix **C**. In this work we exclude all noises except the intrinsic sensor noise. Previous research in the field of yttrium-iron garnet magnetometers [40–42] do not demonstrate any cross-talk phenomenon, while the cross-talk of the most sensitive sensor presented in [23] will be investigated during further studies. In the current paper, since we do not have enough information about any cross-talk, we exclude it from our computations, which makes the sensors independent of each other. In this case, the noise covariance matrix **C**_*YIGM*_ takes the form of diagonal matrix with dispersion of intrinsic noise *σ*_*YIGM*_ at the diagonal. We note that the same reasoning is applicable to other sensors types under consideration of this work. Thus, we assume the matrices **C**_*SQUID*_, **C**_*OPM*_ to be diagonal matrices with values *σ*_*SQUID*_ and *σ*_*OPM*_, respectively. The values of these *σ* are presented below, in Table 1 in Section 4.

**Table 1:**
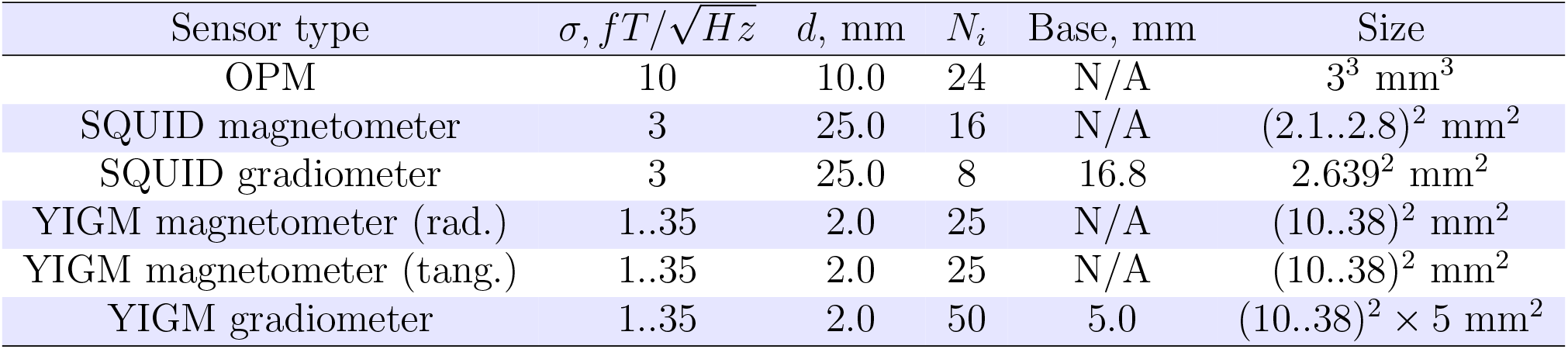
Parameters of all sensors used in this study (from left to right): intrinsic noise *σ*, distance to scalp *d*, number of integration points *N*_*i*_, base and size.

| Sensor type | $\sigma, fT/\sqrt{Hz}$ | $d, \text{ mm}$ | $N_i$ | Base, mm | Size |
| --- | --- | --- | --- | --- | --- |
| OPM | 10 | 10.0 | 24 | N/A | $3^3 \text{ mm}^3$ |
| SQUID magnetometer | 3 | 25.0 | 16 | N/A | $(2.1..2.8)^2 \text{ mm}^2$ |
| SQUID gradiometer | 3 | 25.0 | 8 | 16.8 | $2.639^2 \text{ mm}^2$ |
| YIGM magnetometer (rad.) | 1..35 | 2.0 | 25 | N/A | $(10..38)^2 \text{ mm}^2$ |
| YIGM magnetometer (tang.) | 1..35 | 2.0 | 25 | N/A | $(10..38)^2 \text{ mm}^2$ |
| YIGM gradiometer | 1..35 | 2.0 | 50 | 5.0 | $(10..38)^2 \times 5 \text{ mm}^2$ |

### 3.1 Signal power and signal-to-noise ratio

Since the i^*th*^ column of the matrix **L** defines the output of all *N*_*s*_ sensors with respect to the i^*th*^ unit source location, it represents the magnetic field pattern of the i^*th*^ source, and is called the source topography metric (**t**_*i*_). The source topography allows to estimate the sensitivity in terms of SNR, using the representation (6). Indeed, by introducing the value *q* of source variance for independent sources, the SNR of the i^*th*^ source can be explicitly obtained in the form

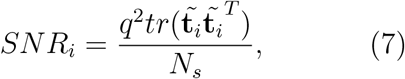

where 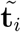 means the whitened source topography: 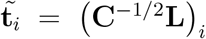. Thus, the SNR metric has the same size as the source space, and therefore allows to obtain convenient brain map in terms of sensitivity of the sensor array with respect to sources located in different areas of the cortex. The *l*_2_-norm of the source topography squared, ||**t**_*i*_||^2^, is defined as the topography power, or signal power. Equation (7) can be simplified due to sensor noise variance *σ*^2^ is equal for all sensors in the array. In this case the topography power is linearly proportional to the SNR of source:

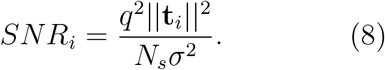

Speaking of relative signal power and relative SNR for two different sensor arrays, they can be calculated in the following way:

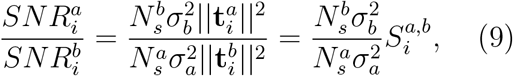

where 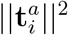 and 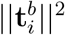 are the topography power of the same source *i* for the arrays *a* and *b*, respectively, and 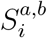 is the relative signal power.

### 3.2 Information capacity

Adapted for MEG/EEG measurements by J.Schneiderman, the total information capacity is related to original Shannon’s channel capacity (see [33]). In the current research, the information capacity was computed in a way proposed in [21], using the orthogonalized SNR of the channels 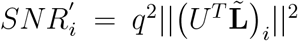. Here the vectors *U* are the eigenvectors of the matrix 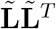. The information capacity can be calculated as follows:

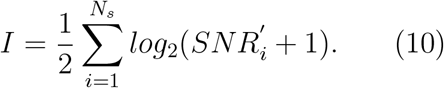

Opposed to the source power and SNR, this metric is an integral and scalar and enable to estimate the effectiveness of the sensor array in terms of the overall information quality gathered in it.

## 4 Sensors

### 4.1 Description of the sensors

In the current paper, we use three types of sensors for our computations: YIGM, OPM, and SQUID. The YIGM sensor is a solid-state magnetometer based on specially shaped yttrium-iron garnet thin films. As it was said earlier, the YIGM sensor is flat by design, having dimensions of 38 × 38 × 3 *mm*^3^. The sensor has two perpendicular sensitive axes, both lying in the sensor’s plane. There are two possible sensor placement according to head: normally to head surface (Figure 1a), i.e. the normal to the head coincides with one of the sensitive axis, and in a tangential way (Figure 1b).

**Figure 1:**
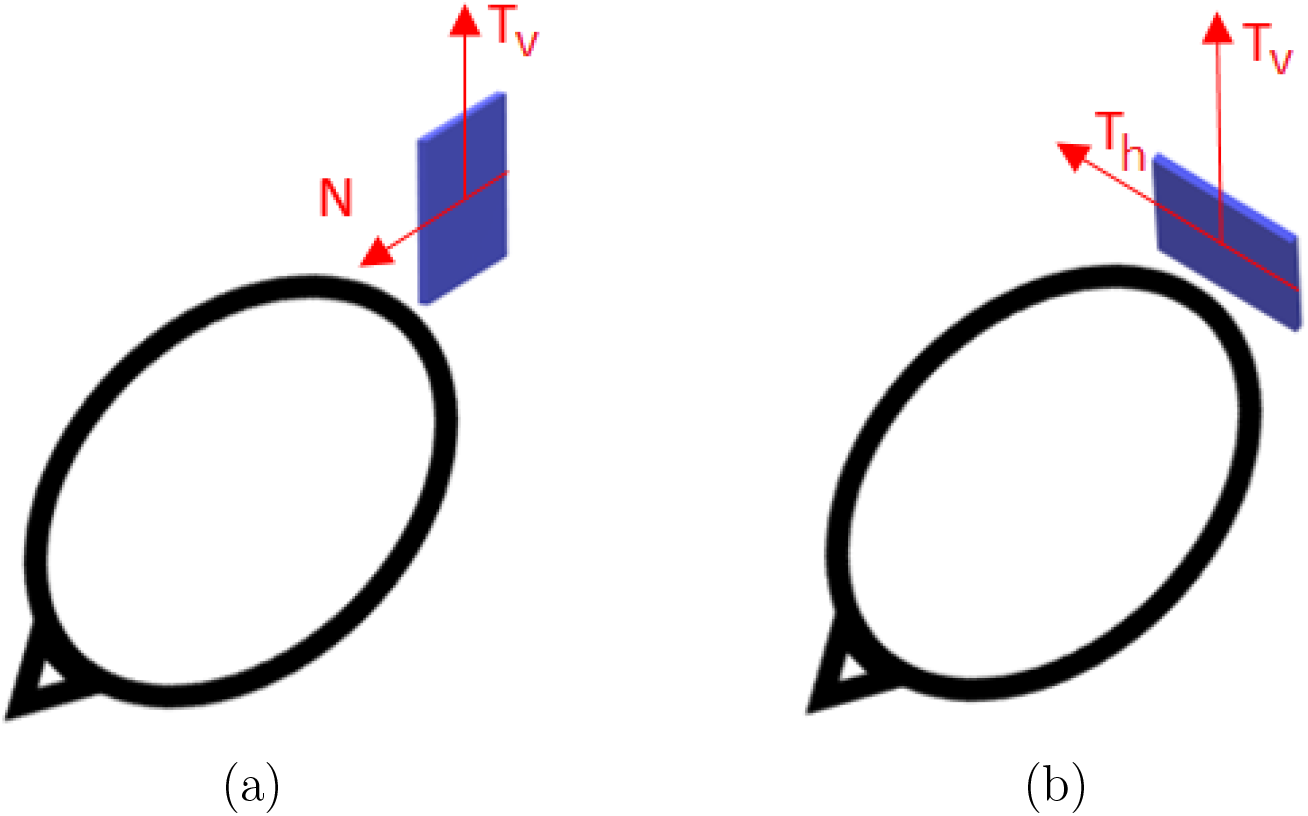
Placement of the YIGM sensor for measurements of radial (a) and tangential (b) components of the magnetic induction.

In the first case, it is possible to register radial (normal) and one of the tangential components of magnetic induction (in the sensor’s plane). In the second case, we can register one or two tangential components equidistant from the head. We focus readers’ attention and stress the fact that here the distance between the nearest housing of the sensor and surface of the head (scalp) can be minimized down to units of millimeter. Since the magnetic field is rapidly decaying, we assume to obtain better SNR in YIGM tangential measurements rather than in normal ones.

We also note that, taking into account the absence of evident cross-talk phenomenon in previous YIGM measurements, it could be possible to construct the gradiometer, using two YIGM sensors. Here the sensors are placed parallel to each other with 5 *mm* sensor-to-sensor distance in the “normal” configuration with respect to the head (see Fig. 2). In this case, the spatial tangential derivative of normal component and one of the tangential components of the magnetic induction vector can be obtained. It is, however, impossible to prove at the moment that this scheme is feasible for measurements of normal derivative of the tangential component. Thus, in this research we use only the scheme of gradiometric measurements presented in Fig. 2. For comparison the same Fig. 2 contains the scheme for gradiometers implemented in reference SQUID system.

**Figure 2:**
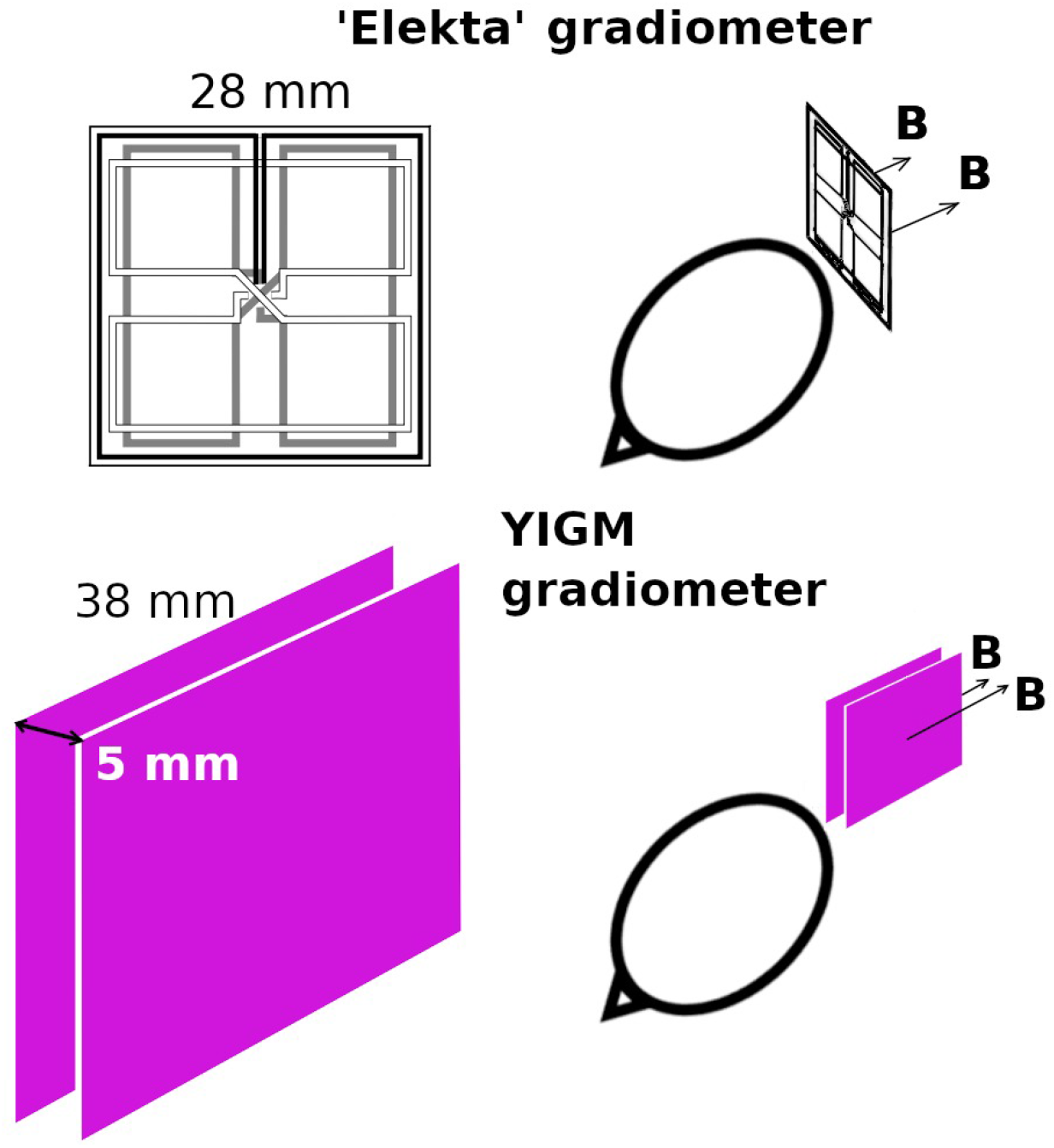
Representation of the gradiometric scheme for measurement of tangential derivative of the radial component of the magnetic induction vector 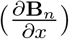 and Elekta’s gradiometric scheme for comparison.

In our study we also use QZFM OPMs (QuSpin Inc., USA) and the SQUID-system Elekta Neuromag (Elekta Oy, Finland). Their parameters are reflected in numerous literature (see, e.g. [7,8,21,27]). The parameters for all three sensor types are compiled in Table 1. Here by size we mean the size of sensor sensitive element, or working body. In case of OPMs it is cubic glass cell inside the sensor itself, while in case of SQUID and YIGM it is a plane. The intervals presented in this column represent the different SQUID machines and YIGM sensors of different size used in this paper (see in details in Subsection 4.2). Correspondingly, by distance to scalp *d* was meant the distance from the center of glass cell in case of OPMs, the distance from the coil plane for SQUID and the distance from the nearest housing in case of YIGM. The 10 mm distance for OPMs is due to the sensors heat problems reported in [13, 36] and to ensure the comfort of the subjects.

### 4.2 Output computation

For each sensor type the sensor output was obtained by the approximate integration over the sensor’s working body, using the set of certain integration points within it. Since both SQUID and OPM sensors are considered to be well-established for MEG purposes, for these sensor types the integration points (as well as some other parameters) are fixed in different packages related to MEG/EEG. In the current research, the forward problem solution (the lead-field matrix) was computed in MNE-Python [1], using the points listed in the file ”coildef.dat”. For short, we do not list the coordinates of these points here, stressing only the number of them (shown in the fourth column of Table 1).

Taking into account the linear dimensions of the zero-generation of the YIGM sensor, the magnetic field could differ a lot over the sensor’s working body (plane), especially in normal component measurements (see Fig. 1, a). To decrease the integration error we apply rather dense mesh for integration: 5 × 5 points, *N*_*i*_ = 25. The nodes of this mesh are distributed uniformly over the integration plane. The coordinates of integration points therefore depend on linear size of the sensor. In this paper we test four different possible sizes. The current one is 38 × 38 *mm*^2^, while shortly we are planning to decrease it to 30 × 30 *mm*^2^ or 20 × 20 *mm*^2^. We also consider the theoretically possible size of 10×10 mm for the future. Since the YIGM sensor is novel the information about its integration points was added to the ”coildef” file in the standard manner. Again, for brevity, we do not list the coordinates of these integration points here.

Sensor’s coordinates and orientations were stored in mne.info structure for each sensor type and each layout listed below in Subsection 4.3.

### 4.3 Sensor arrays

To compare different layouts of sensors, we choose two schemes of sensor locations. The first one is taken from existing and well-established MEG system Elekta Neuromag (further we refer to it as ’SQUID-like layout’). The second one is defined using uniform distribution of sensors over the scalp (area of measurement).

#### 4.3.1 SQUID-like layout

To obtain the layout implemented in Elekta system we first apply typical digitization. We then define the point on the scalp, such that the line connecting it with the center of the SQUID sensor is aligned with the normal to the head surface (see Fig. 3). After that we place OPM and YIGM sensors on this line at a specific for each sensor distance stated in Table 1. This procedure is repeated for the rest of the points obtained on the stage of digitization.

**Figure 3:**
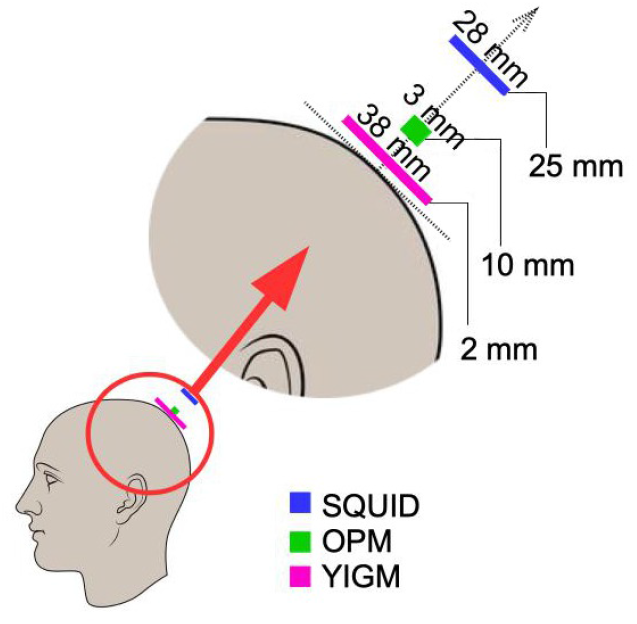
Positions of sensors of different types with respect to head surface.

In this manner, we obtained three SQUID and OPM layouts used for further computations together with a set of YIGM layouts. Original SQUID system provides us with two SQUID layouts, namely: containing 102 magnetometers (see Fig. 4, a) and 204 planar gradiometers (not shown). For OPMs we obtained the SQUID-like layout consisting of 102 points that are locations for centers of OPM gas cells (Fig. 4, b). For YIGM we formed three layouts: one consisting of 102 normally-oriented YIGMs (Fig. 4, c), one for tangential component measurements (102 tangentially-oriented YIGMs), see Fig. 4, d), and the last one for gradiometric measurements of the radial component (see Fig. 4, e).

**Figure 4:**
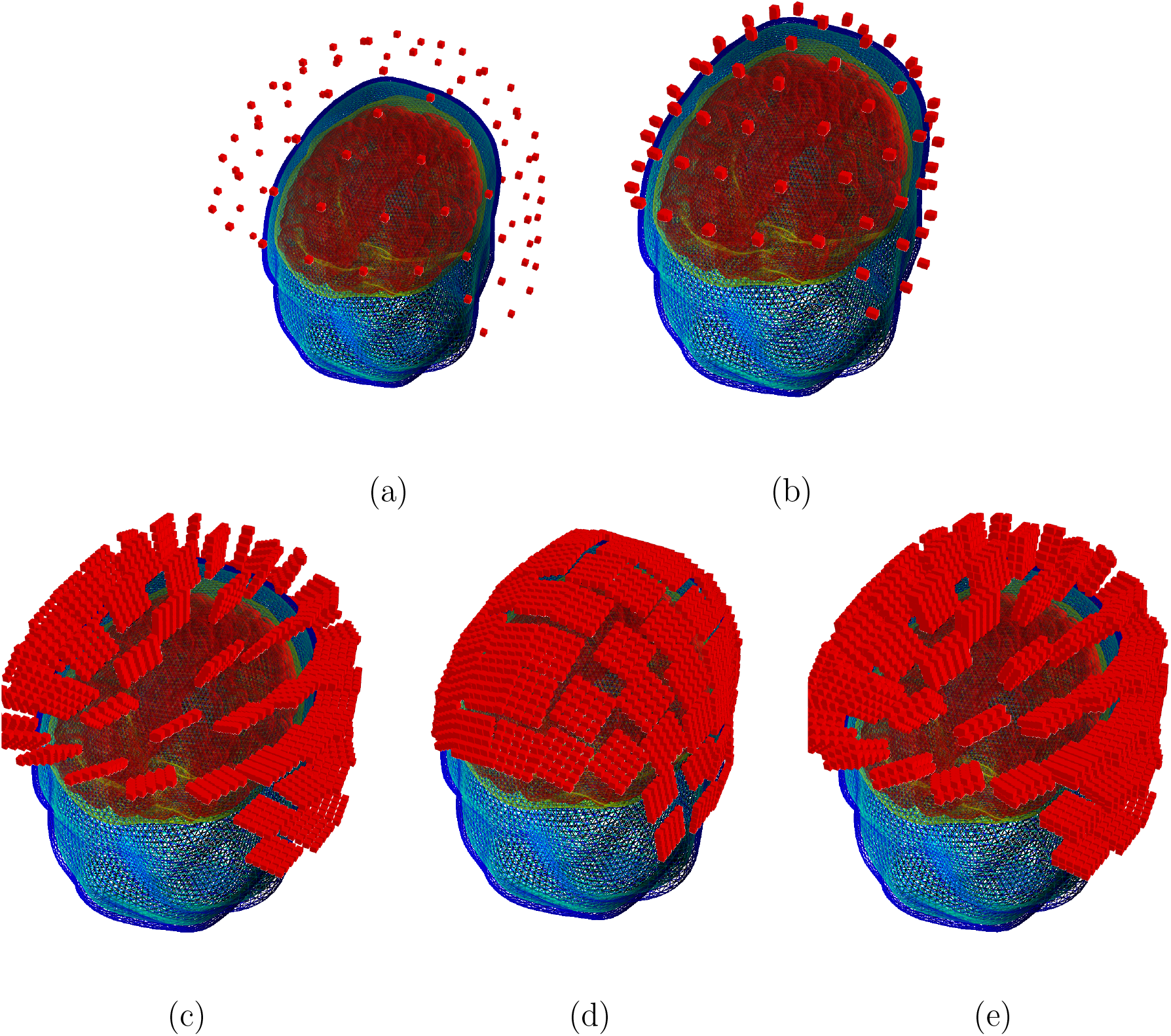
SQUID-like layouts: (a) - original SQUID layout (Elekta Neuromag MEG system); (b) - OPM SQUID-based layout; (c) - YIGM-NL (normally-oriented) SQUID-based layout; (d) YIGM-TG (tangentially-oriented) SQUID-based layout; (e) - YIGG-NL (gradiometric scheme with sensors normally-oriented) SQUID-based layout.

We use 2 mm distance from scalp to the nearest edge or plane of the YIGM sensor. Speaking of the distance between the scalp and the center of the sensor plane, it is the same 2 mm for tangential measurements (we use the sensor size of 20 × 20 mm to avoid intersections). In case of normally-oriented sensors it depends on size of the YIGM sensor as *d* = 2.0 + *size/*2 (mm). Since we worked with squares with a side of 10, 20, 30, and 38 mm, these distances are 7, 12, 17, and 21 mm, respectively. For brevity, in Fig. 4 we show only YIGM layouts generated for YIGM plane of 20 × 20 mm size.

#### 4.3.2 Uniform layouts

Another kind of sensor layout used in the current study was created originally for our implementation of the OP-MEG system [23]. In this layout, sensors are distributed uniformly over the area of measurements (scalp). The coordinates were chosen in such a way that YIGM sensors, placed for tangential measurements, do not intersect with each other. Therefore, the number of sensors in layout depends on the sensor size. Thus, here we explore not one layout, but a set of layouts, main parameters of which are shown in the Table 2. As with SQUID-like layout, we used 2 mm distance from scalp to the sensor’s edge for YIGMs. Sensors together with their integration points for these layouts are presented in Fig. 5.

**Table 2:** Uniform layouts: number of sensors *N*_*s*_ depending on the sensors’ sizes. OPMs’ sizes are not shown since they are constant.

| Size, mm<sup>2</sup> | 38<sup>2</sup> | 30<sup>2</sup> | 20<sup>2</sup> | 10<sup>2</sup> |
| --- | --- | --- | --- | --- |
| $N_s$ | 35 | 45 | 105 | 333 |

**Figure 5:**
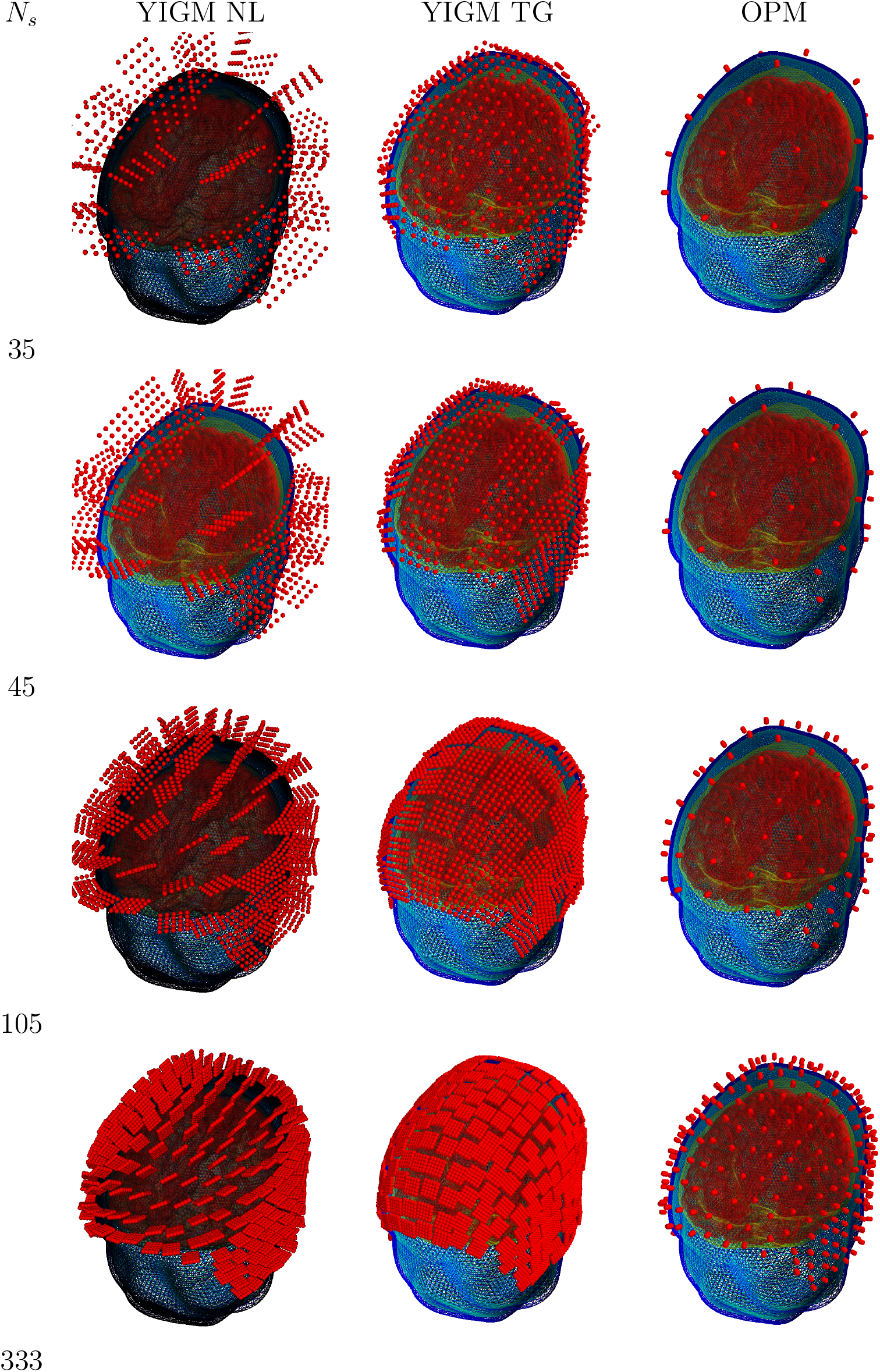
Uniform layouts corresponding to Tab. 2. Number of sensors for each layout triplet (row) is shown on the left side.

## 5 Results

In order to apply metrics mentioned in Section 3, we compute the lead-field matrices for all layouts listed in Section 4.3.

The source power for each layout was computed using the column-wise norm of the lead-field matrix, while the SNRs were calculated via formula 7. For conventional Elekta Neuromag MEG system the source power and SNR were computed separately for magnetometers and gradiometers. In case of YIGM, we calculated the SNRs for noise varying from 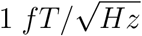, which is the theoretical limit, and 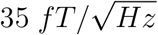, which is the current noise level of YIGMs obtained in [23].

Each layout of both OPM and YIGM was compared to reference Elekta Neuromag layout using representation 9, forming relative signal powers and relative SNRs. Having different noise levels for YIGMs, we formed mean relative powers and mean SNRs.

We also computed the channel information capacity for all layouts using the formula 10. Again, in case of YIGM it depends on the noise level for each layout.

### 5.1 SQUID-like layouts

Firstly, we explore the SQUID-based layouts described in Section 4.3. The dependencies of relative power, relative SNR, and channel information capacity are shown in Fig. 6. These computations have been done using the noise levels presented in Table 1. In order to make the plots more informative, we used logarithmic scale along the vertical axis.

**Figure 6:**
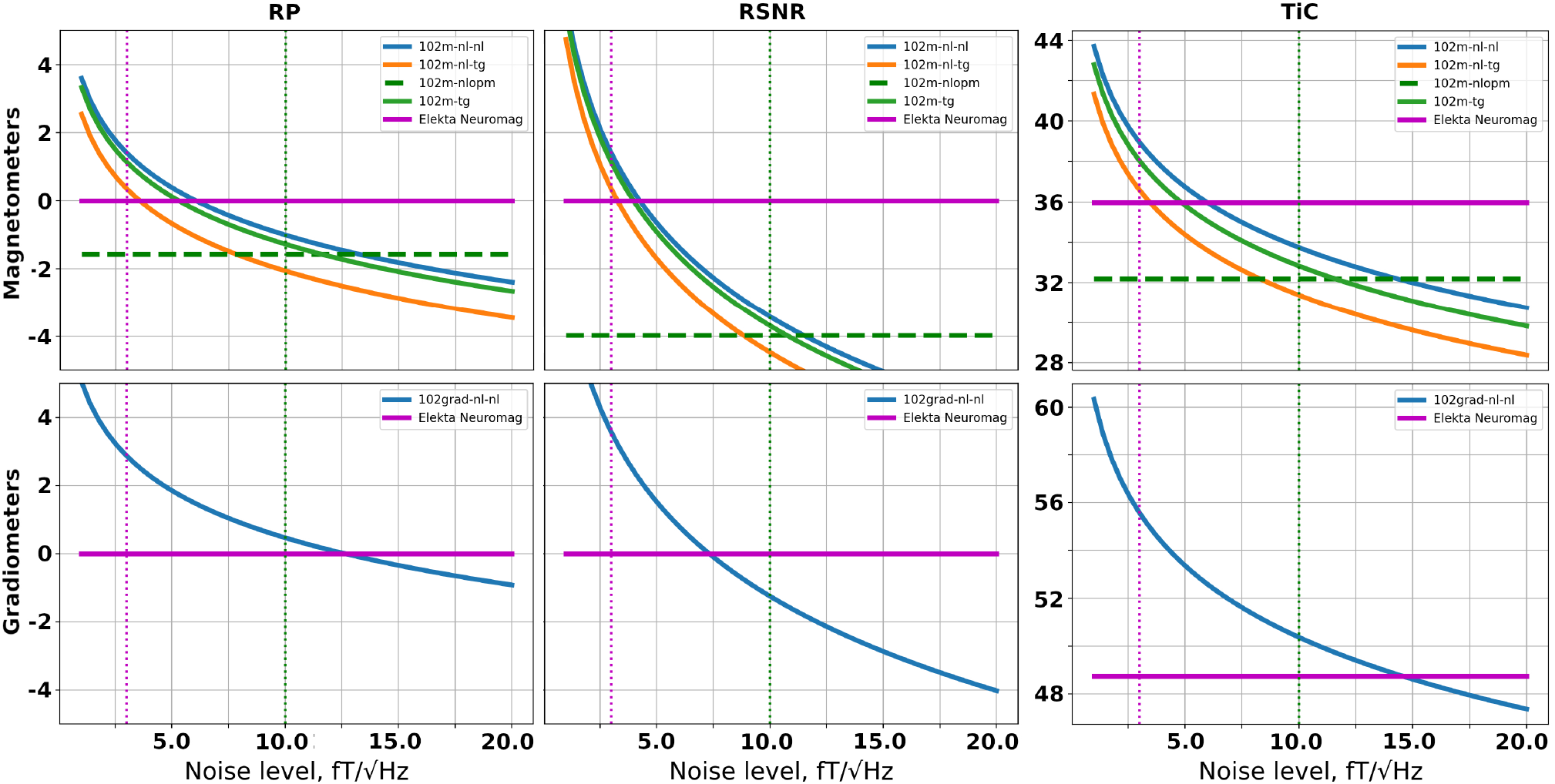
Dependencies of relative signal power (left), relative SNR (middle), and total information capacity (right) for SQUID-based layouts formed with: magnetometers (top row), gradiometers (bottom row).

The top row shows metrics for SQUID-based layouts consisting of magnetometers. The performance of the original Elekta Neuromag layout shown with solid magenta line, the performance of the SQUID-based layout formed by OPMs is presented with dashed green line, while solid blue, orange and green lines are responsible for YIGM layouts. Please note in the legend that YIGM layouts are labeled by orientation of the sensor first and then followed up by the component measured. In case of tangential orientation of YIGMs both components showed approximately the same results therefore we left only one of them. Since metrics for Elekta and OPMs were calculated using constant noise levels they represent constants as well. Dotted magenta and green vertical lines correspond to the noise level of Elekta 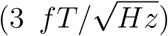 and OPMs 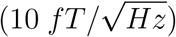, respectively, and added for convenience.

Among YIGM layouts, normally-oriented sensors measuring normal component show the best results, while tangentially-oriented YIGMs reveal lower metrics values. The lowest metrics is due to normally-oriented sensors measuring tangential component. YIGM layouts show better results than SQUIDs in case of 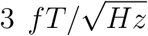 noise level and than OPMs in case of 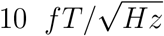 noise level in all three metrics. However, to outperform the conventional SQUID-based MEG system Elekta Neuromag consisting of 102 magnetometers we need the intrinsic noise level of YIGMs to be 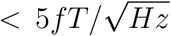, and 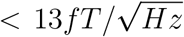 to outperform the OP-MEG developed within the same montage.

The bottom row in Fig. 6 shows the same for gradiometers, taking into account the fact that only one type of YIGG was used: gradiometers registering tangential gradient of the normal component and oriented normally (presented by solid blue line). The relative signal power, relative SNR and channel capacity of YIGGs are compared only with that of original Elekta’s gradiometric layout (204 planar gradiometers) shown in bold magenta line since we do not consider gradiometers based on OPMs in this paper. The gradiometric scheme with assumed noise level 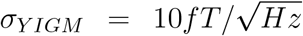 outperforms both SQUID gradiometers and OPMs in terms of relative signal power and channel capacity, but shows lower SNR. In case of 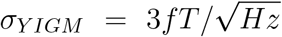 the YIGG SQUID-like layout dramatically outperforms both SQUIDs gradiometers and OPMs in terms of all metrics. However, to achieve the same SNR SQUID gradiometers do the noise level of YIGG must be 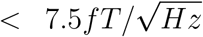. At 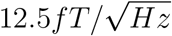 and 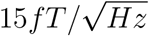 it starts to outperform Elekta’s gradiometric layout in terms of relative signal power and total information capacity, respectively.

### 5.2 Uniform layouts

In this section, we compare the uniform layouts formed with YIGMs and described in Section 4.3 with conventional SQUID-MEG system (Elekta Neuromag) as well as between each other. Since these layouts have different number of sensors, we can explore how the required noise level changes depending on the different number of sensors. As before, in order to compute relative signal powers and relative SNRs we use the Elekta Neuromag as reference system.

The dependencies of metrics on noise level for different uniform layouts are shown in the Fig. 7. The structure of this figure is similar to that of Fig. 6. With increase in number of sensors in a layout the metrics value grows for both magnetometer types and for gradiometers as well.

**Figure 7:**
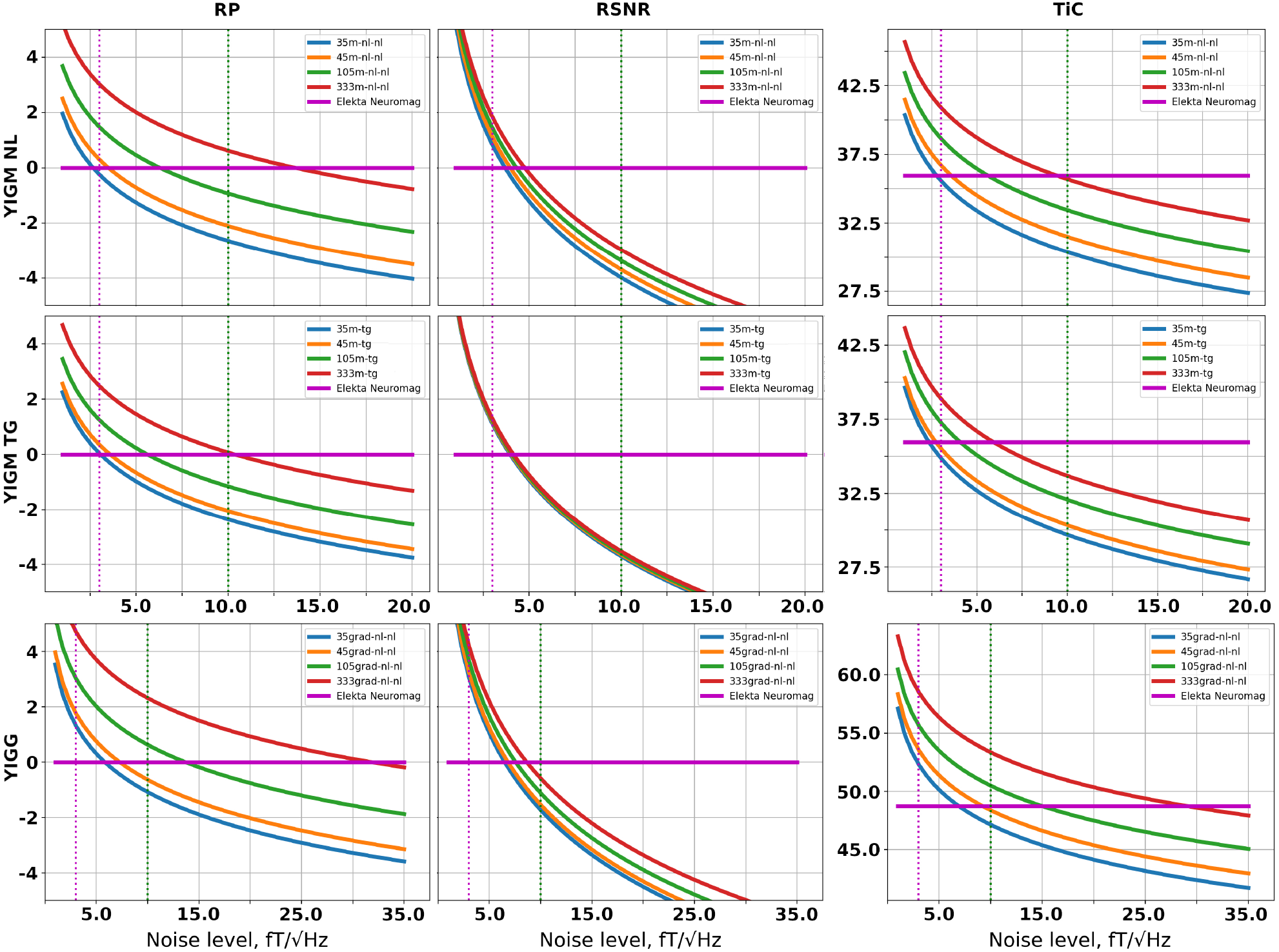
Relative power (left column), relative SNR (middle column), and total information capacities (right column) computed for different uniform layouts of YIGM: magnetometers (first and second rows), and gradiometers (bottom row).

Considering the 333m-nl-nl layout, we start to outperform Elekta Neuromag at level of 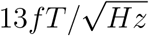 and 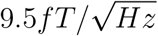 in terms of signal power and information capacity, respectively. As for SNR, the noise level required is still about 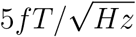. Like for SQUID-like layouts the uniform layout with YIGMs located tangentially show lower metrics compared to normally-orientated. Moreover, it is interesting to note that for tangential layouts difference in RSNR is subtle and we see all four curves coinciding.

The 333grad-nl-nl layout allows to outperform Elekta’s gradiometers at the noise level of about 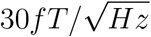 in terms of relative signal power and total information capacity. The numbers for outperforming Elekta in SNR are about at least 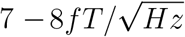.

Figure 8 shows the visual representation of required number of channels in two types of magnetometric as well as gradiometric layouts for certain intrinsic noise level of YIGM sensors. The red line indicates the level of q = 1, i.e the same value as for Elekta Neuromag in terms of signal power (left panel) and information capacity (right panel). The dependencies for relative SNR represent almost straight perpendicular vertical lines, i.e. no dependency between intrinsic noise and number of channels in layout was detected. For brevity the corresponding figures are not shown.

**Figure 8:**
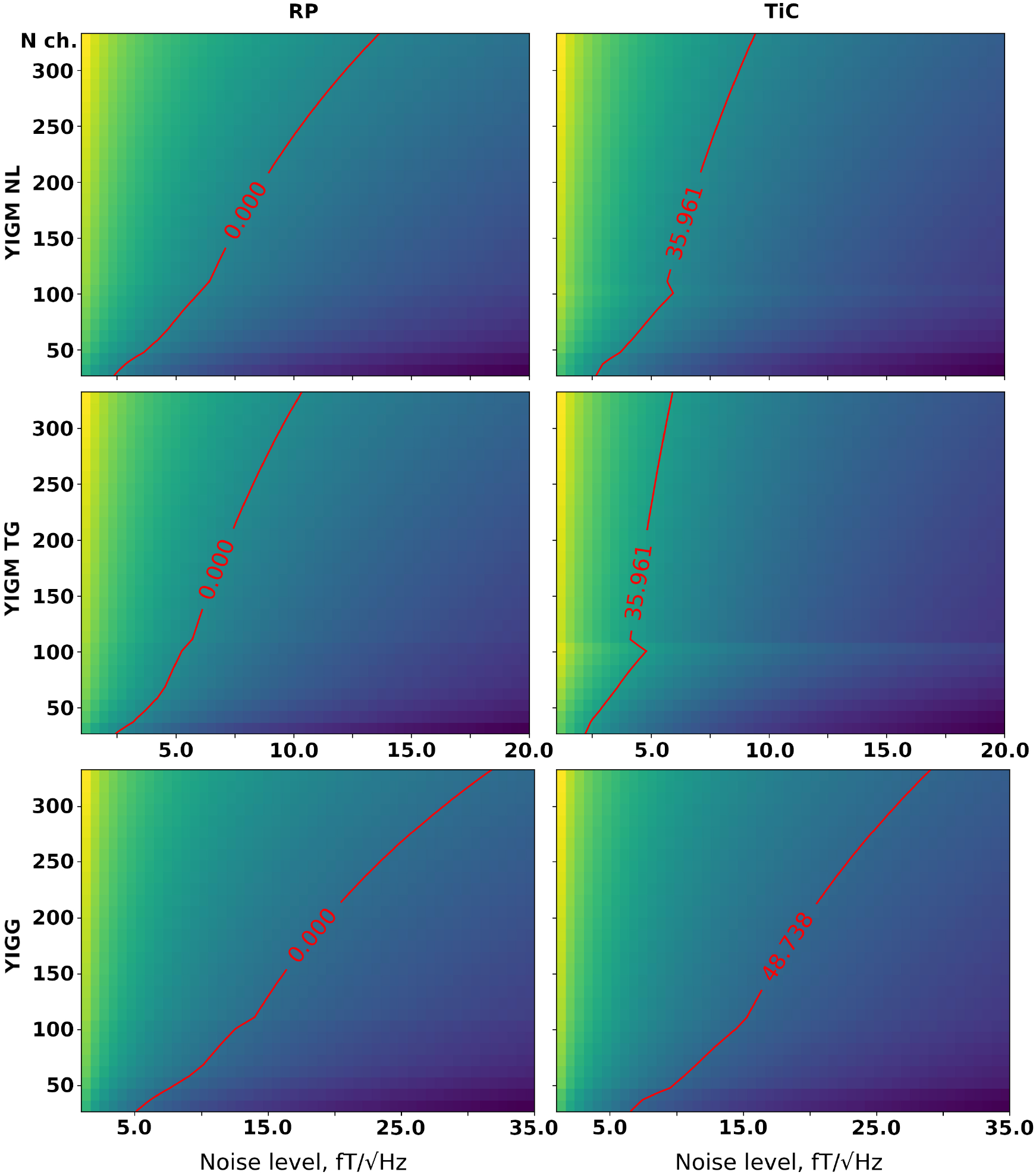
Dependence of relative signal power (left column), and total information capacity (right column) on the number of channels and on different levels of intrinsic noise compared to conventional Elekta Neuromag SQUI-MEG system. The top row represents normally-oriented YIGMs, the middle row - tangentially-oriented YIGMs, and the bottom row shows the metrics for normally-oriented YIGM-based gradiometers with the base of 5 mm. The red line represents the value q = 1.0, meaning the equal metrics to that of reference SQUID-MEG system Elekta Neuromag.

For additional motivation, to investigate the power of coverage we show relative YIGM/OPM and relative YIGM/Elekta SNR cortical maps for magnetometers in Figure 9. We use three layouts:, minimal uniform layout (35 sensors), Elekta-like layout (102 magnetometers) and maximum uniform layout (333 sensors). Since normally-oriented YIGMs measuring normal component perform better, we use only them for these calculations. Noise level of YIGMs was chosen in such a way to be equal to that of comparing type of magnetometer - OPM or SQUID - 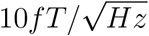 and 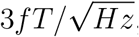, respectively. The scale is from 0.0 to 10.0, where blue color means bad coverage, red - good coverage, and green is responsible for the equal coverage as for comparing type of magnetometer. Increase in number of channels gives better coverage, moreover YIGM vs. OPM coverage give considerably different result.

**Figure 9:**
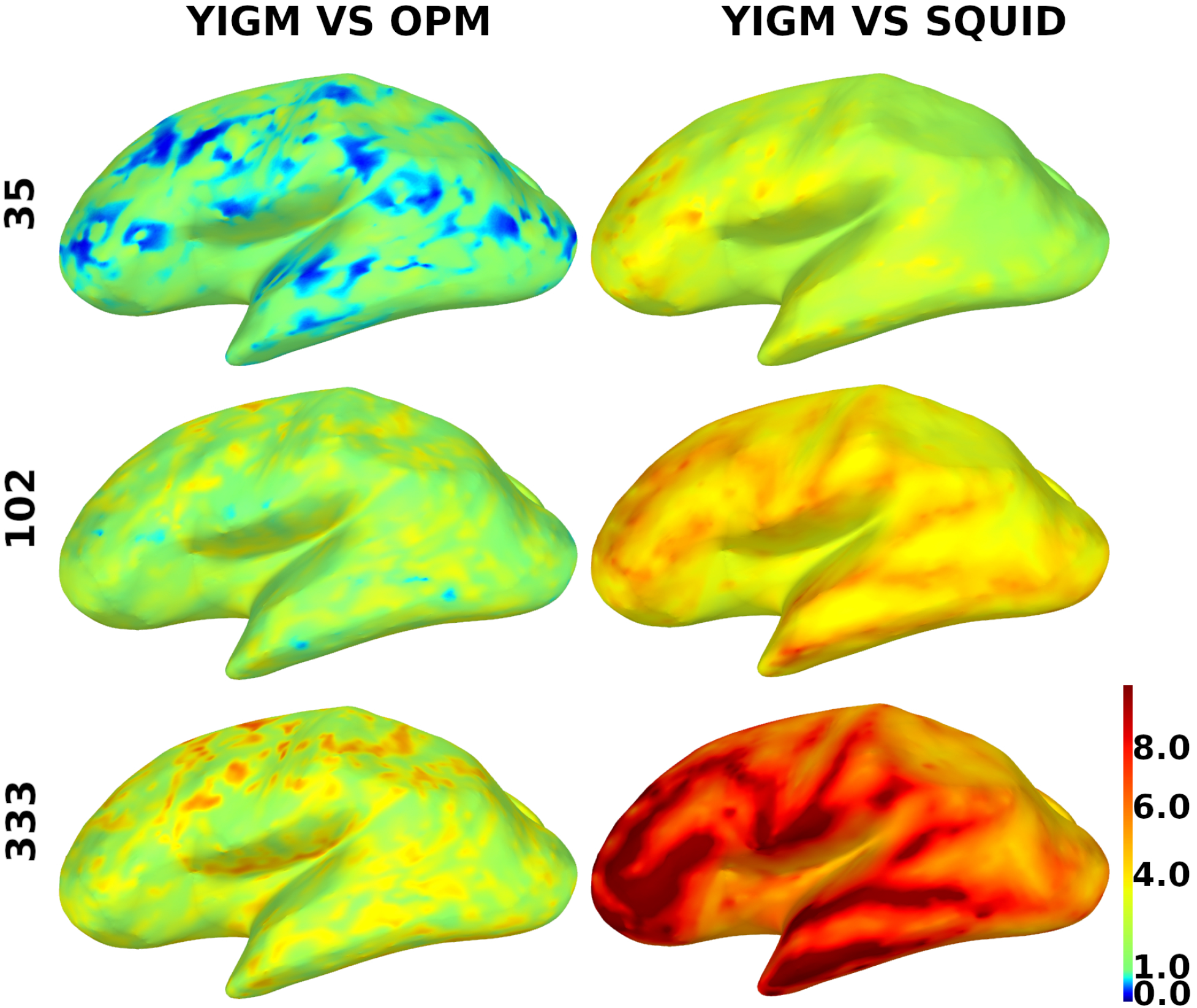
Distribution of the relative SNR over the cortex for magnetometers. YIGM layouts (minimum uniform, SQUID-like and maximum uniform) with noise level 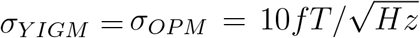 compared with OPMs (left panel) and with noise level 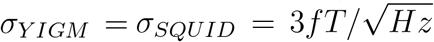 compared with SQUID gradiometers (right panel). The scale is from 0.0 to 10.0.

Fig. 10 shows the same for gradiometers, except the scale. Since gradiometers are a fundamentally different thing the scale here is from 0.5 to 60.

**Figure 10:**
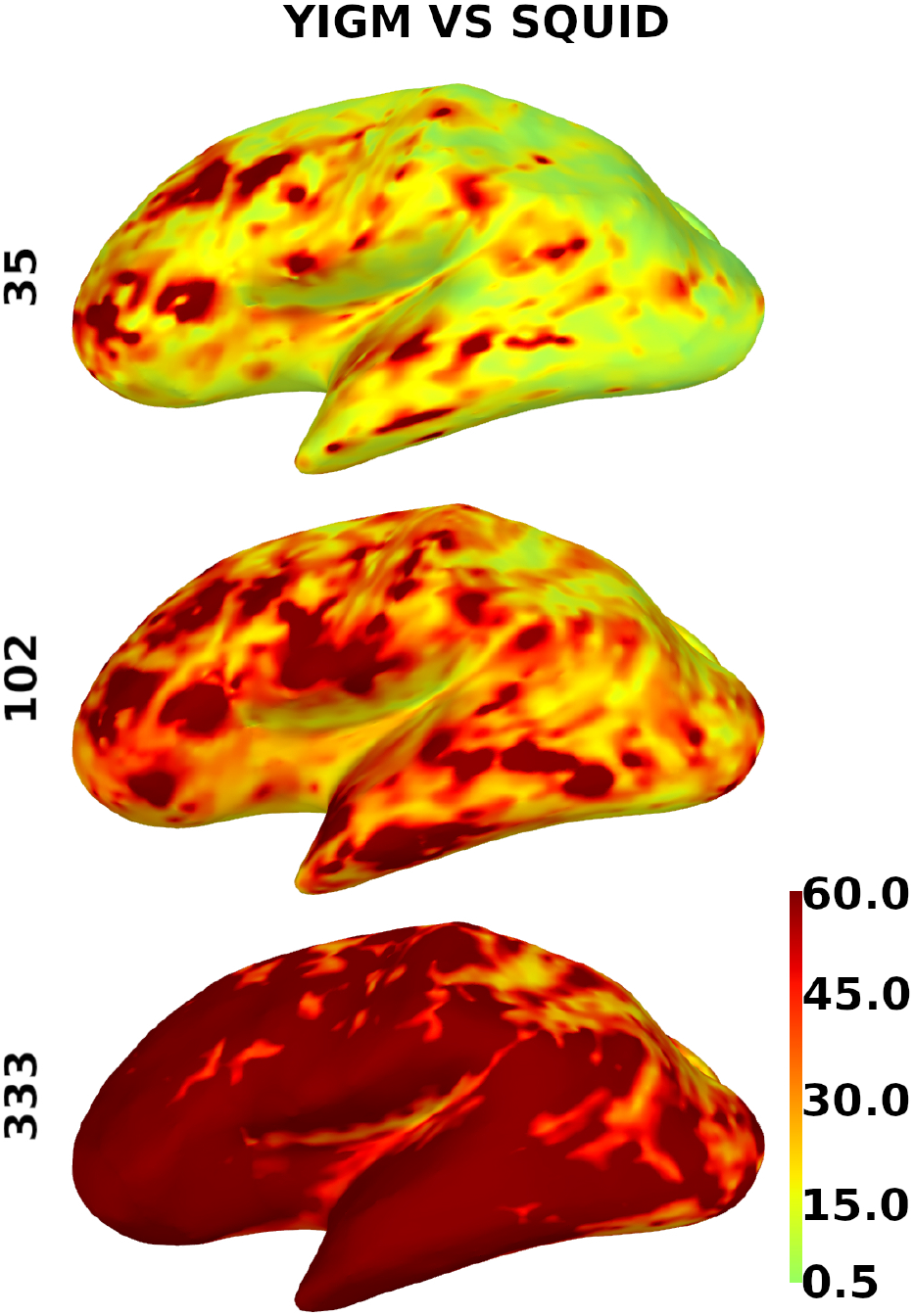
Distribution of the relative SNR over the cortex for gradiometers. YIGM layouts (minimum uniform, SQUID-like and maximum uniform) all with noise level 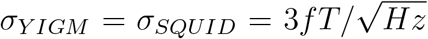 compared with SQUID gradiometers. The scale is from 0.5 to 60.

## 6 Discussion

The results presented above may give some hints for further development of multichannel YIG-MEG system. Here we present some qualitative analysis of the quantitative results.

Analyzing Figure 6, we conclude that the current noise level of YIGM sensor 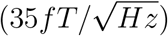 is unfortunately insufficient for construction of the multichannel YIG-MEG system. Although sensors are closer to the head surface and therefore outperform SQUID and OPMs at the level of their noise levels, the overall SNR of the MEG system built with usage of such magnetometers is rather low. Weak dependence of the overall SNR of sensor array on the number of channels described above with figure 8 is expected taking into account the assumption of independence of the sensors in an array with respect to each other.

We can conclude that in case of the most dense layout consisting of 333 magnetometers, we need the YIGM noise level to be already two times less than the current one to outperform Elekta Neuromag in terms of signal power. However, as we stated above, the SNR is not affected significantly by number of sensors, and the noise level required to outperform Elekta Neuromag in terms of SNR is still far from what we have.

The values of the computed metrics seem a bit better for sensor layouts compiled from gradiometers. Indeed, it is expected that for cortical sources gradiometers perform better than magnetometers. Figures 9 and 10 with cortical maps confirm this, since gradiometers provide better coverage. However, again, it is obvious that the current intrinsic noise level of YIGM is still far away from SQUID’s numbers.

Remarkably that in case of the most dense uniform YIGG layout (333 gradiometers) the YIGG noise level required for outperforming Elekta in relative signal power and total information capacity is very close to the current YIGM noise level, but even here, without exception, SNR is a stumbling block.

Speaking of YIGM tangential layouts, as was said earlier, we obtain considerably poorer results compared to normal ones. At first thought, it seems to be illogical, since in the case of tangential placement all points on the plane of YIGM sensor is more or less equidistant from the scalp surface. However, as was stated in Subsection 1.2 weakness of tangential layouts can be explained by the fact that the magnetic field caused by volumetric currents substantially suppresses the overall amplitude of the topography created by primary currents in tangential measurements [21]. In this way, the decrease in SNR becomes plausible. Indeed, a separate study is needed to find out more.

Short comment on YIGM vs. OPM cortical maps. Since the comparison between YIGM and OPM was a secondary goal of this paper, we did not devote much attention to this, however, it is clear from the Figure 9 that with the same noise level, the YIGM does not give a considerable advantage in coverage even with the most dense layout. The completely different picture is observed in YIGM vs. SQUID maps, where 333 YIGM sensors provide brilliant coverage, especially in frontal and temporal regions. Of course, an additional research is required to establish the cause of this difference.

Despite all above, it is important to note that in case of approaching the theoretical limit for YIGM sensors 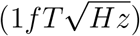, the YIG-MEG system with 102 either gradiometers or magnetometers may outperform Elekta Neuromag in 20 − 60 times in terms of SNR and signal power, and by 15−40% in terms of channel capacity. Since the approach to the theoretical sensitivity for YIGMs is an engineering versus scientific issue, we assume these values to be a good motivation for further YIGM/YIGG development.

## 7 Conclusions

In this simulation study we explored different possible layouts for multi-channel YIG-MEG systems and via the lead-field analysis compared the performance of YIGMs with conventional SQUID system as well as well-established OPMs. The values of relative source power, relative SNR and total information capacity revealed that the primary step towards efficient multi-channel YIG-MEG system is to reduce the YIGM sensor noise, since the increase in the number of channels (decrease in size) with current intrinsic noise of 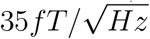 does not give a visible improvement in YIGMs performance.

Our future work on forward modeling will be devoted to calculation of optimal locations for sensors on the head, using more complex principles than just uniform distribution, as well as the possible usage of an averaged head model instead of individual one. Moreover, in this paper we used MNE, which in its turn involves the boundary-element method (BEM) for MEG modeling. Therefore, there is an idea to try different methods. We refer to [43] for a discussion on the BEM as well as standard and more advanced schemes, such as finite-element method (FEM). Finally, special attention should be paid to tangential YIGM layouts and their effectiveness.

